# A robust *ex vivo* experimental platform for molecular-genetic dissection of adult human neocortical cell types and circuits

**DOI:** 10.1101/325589

**Authors:** Jonathan T. Ting, Brian Kalmbach, Peter Chong, Rebecca de Frates, C. Dirk Keene, Ryder P. Gwinn, Charles Cobbs, Charles Cobbs, Andrew L. Ko, Jeffrey G. Ojemann, Richard G. Ellenbogen, Christof Koch, Ed Lein

## Abstract

The powerful suite of available genetic tools is driving tremendous progress in understanding mouse brain cell types and circuits. However, the degree of conservation in human remains largely unknown in large part due to the lack of such tools and healthy tissue preparations. To close this gap, we describe a robust and stable adult human neurosurgically-derived *ex vivo* acute and cultured neocortical brain slice system optimized for rapid molecular-genetic manipulation. Surprisingly, acute human brain slices exhibited exceptional viability, and neuronal intrinsic membrane properties could be assayed for at least three days. Maintaining adult human slices in culture under sterile conditions further enabled the application of viral tools to drive rapid expression of exogenous transgenes. Widespread neuron-specific labeling was achieved as early as two days post infection with HSV-1 vectors, with virally-transduced neurons exhibiting membrane properties largely comparable to uninfected neurons over this short timeframe. Finally, we demonstrate the suitability of this culture paradigm for optical manipulation and monitoring of neuronal activity using genetically encoded probes, opening a path for applying modern molecular-genetic tools to study human brain circuit function.

## Introduction

There is a tacit assumption in much of modern neuroscience that brain structure and function are largely conserved between rodents and humans. However, the limited success in translating findings from rodent models of disease into effective new treatments for human disease implores us to re-examine this assumption^1,2^. In this regard, direct measurement of the functional properties of the human brain at the resolution of cell types and circuits is invaluable and absolutely necessary to delineate what features are conserved or not relative to the ubiquitously studied rodent brain.

One promising avenue to advance our knowledge on this topic has been the use of *ex vivo* brain slices derived from adult human neurosurgically-excised tissue to probe the structure and function of living human neurons and synapses by applying standard procedures originally optimized for rodent neurophysiology research^3–5^. A number of differences have been described between the human and rodent brain in support of functional divergence, including distinctive synaptic transmission and plasticity mechanisms ^6–8^, differential effects of neuromodulators ^9^, unique cell types ^10,11^, and provocative evidence of a lower specific membrane capacitance of human pyramidal neurons ^12^. Collectively, these findings highlight some overt and important limitations of animal model research for understanding the human brain. Thus, more functional studies on human *ex vivo* brain specimens are warranted in order to provide a critical knowledge base to better guide translational research towards more effective treatments for a wide range of human neurological and psychiatric disorders.

Efforts to diversify and extend the uses of adult human *ex vivo* brain slices could prove transformative, particularly given that availability of this precious biological material is limited and likely declining with increasing prevalence of less invasive neurosurgical techniques ^13^. One notable limitation of human *ex vivo* brain slice studies to date has been the inability to leverage molecular genetic tools to prospectively mark, monitor, and manipulate cell types to more precisely explore their functional properties and contributions within brain circuits. In contrast, the extensive and diverse repertoire of molecular genetic tools and strategies applicable to the mouse has enabled and accelerated new insights into the role of specific cell types within brain circuits, functional connectivity, and the circuitry basis of behavior ^14,15^. Earlier efforts to achieve genetic labeling in human brain slices showed limited promise due to poor tissue viability ^16,17^, or limited evidence to demonstrate functional integrity of transduced neurons ^18^. However, more recent efforts have achieved prolonged culture of adult human *ex vivo* brain slices that retain intact electrophysiological function ^19,20^ and, in a separate study, a first demonstration of viral genetic labeling for optical control of human neuron firing using lentivirus to express Channelrhodopsin-2 (ChR2)^21^. This notable breakthrough suggests that systematic genetic targeting and functional dissection of human brain cell types and circuits may be more tractable than previously thought, at least for those brain regions removed during surgical procedures and deemed suitable for research purposes.

Despite these recent advances, most of the evidence has been largely qualitative, with only limited examples provided. As such, important questions remain about the use of cultured and virally-transduced adult human brain slices. How robust are these cultures, and how well preserved are structure and function relative to the gold-standard acute brain slice? The answers to these questions are crucial to understanding the validity, utility, and limitations of the system for deriving new knowledge about human brain cell types and circuit function. Here we adapt a brain slice protocol originally optimized for mature adult rodent brain tissue ^22^ and provide direct quantitative evidence substantiating the robustness and exceptional viability of adult human *ex vivo* brain slices maintained *in vitro*, including evidence that layer organization and neuronal function are well-preserved over early culture times. We also demonstrate rapid virus-mediated transgene expression using Herpes Simplex Virus type 1 (HSV-1) amplicon vectors ^23^ and show the feasibility of marking, monitoring and manipulating human neurons using genetically-encoded probes within this short time window.

## Materials and Methods

### Human surgical specimens

Surgical specimens were obtained from local hospitals in collaboration with a network of neurosurgeons. All patients provided informed consent for tissue donation, and all experimental uses were approved by the respective hospital Institutional Review Board before commencing the study (the Swedish Institutional Review Board for patients at Swedish Neuroscience Institute or the Institutional Review Board of the University of Washington for patients at Harborview Medical Center). All experiments and methods were performed in accordance with the relevant guidelines and regulations. The patient cohort included both temporal lobe epilepsy and deep brain tumor patients. This study covers 25 surgical cases (19 epilepsy and 6 tumor resections), with an average patient age of 39.9 years old and a range from 19-75 years old. Of these 25 patients, 9 are male and 16 are female. Twenty-one specimens were derived from the temporal lobe, 3 specimens from the frontal lobe, and 1 specimen from the parietal lobe. Neocortical tissue was surgically excised to gain access to deeper pathological brain tissue targeted for resection. The resected neocortical brain tissue was distal to the core pathological tissue and thus deemed suitable for research purposes and not of diagnostic value.
In one case, we obtained hippocampal tissue in addition to the temporal lobe tissue from a temporal lobe epilepsy patient. Every effort was made to ensure clean scalpel cuts at the margins of the resected tissue and to avoid electrocautery unless deemed absolutely necessary. To the extent possible, the scalpel incisions were perpendicular to the pial surface with the intent to avoid crude transection of neuronal morphology across the cortical depth. The typical size of neocortical excisions from epilepsy surgeries is approximately 1cm^3^, which, after accounting for trimming steps and reserving adequate material for post-hoc histological assessments, can yield approximately 20 viable slices for electrophysiology. Tumor surgery-derived specimens are far more variable in size, but most always are smaller as compared to excisions from epilepsy surgeries. Surgically excised specimens were immediately placed in a sterile container filled with N-methyl-D-glucamine (NMDG) substituted artificial cerebrospinal fluid (aCSF) of the following composition (in mM): 92 NMDG, 2.5 KCl, 1.25 NaH2PO4, 30 NaHCO3, 20 4-(2-hydroxyethyl)-1-piperazineethanesulfonic acid (HEPES), 25 glucose, 2 thiourea, 5 Na-ascorbate, 3 Na-pyruvate, 0.5 CaCl_2_·4H_2_0 and 10 MgSO_4_·7H_2_O. The pH of the NMDG aCSF was titrated pH to 7.3-7.4 with concentrated hydrochloric acid and the osmolality was 300-305 mOsmoles/Kg. The solution was pre-chilled to 2-4°C and thoroughly bubbled with carbogen (95% O_2_/5% CO_2_) gas prior to collection. The tissue was quickly transported from the operating room to the laboratory site while maintaining constant carbogenation throughout. Transport time from the operating room to the laboratory was between 14-40 min.

### Acute human ex vivo brain slice preparation

The surgical specimens were closely examined to determine the optimal strategy for blocking and mounting on the vibrating microtome platform. Tissue blocks were trimmed and mounted such that the angle of slicing was perpendicular to the pial surface. This ensures that the dendrites of large pyramidal neurons are preserved relatively intact. No effort was made to remove the pia mater, as this procedure greatly prolongs the processing time and risks damage to the underlying grey matter. Slices of 350 μm thickness were prepared on a Compresstome VF-200 slicer machine (Precisionary Instruments) using the NMDG protective recovery method^22,24^ and either a zirconium ceramic injector blade (EF-INZ10, Cadence) or a sapphire knife (93060, Electron Microscopy Sciences). The slicing solution was NMDG aCSF as above for transport. The slices were transferred into a recovery chamber filled with NMDG aCSF at 32-34°C and continuously bubbled with carbogen gas. After a 12 min recovery incubation, the slices were transferred into a Brain Slice Keeper-4 holding chamber (Automate Scientific; see also ^22^ for more details on chamber design) containing, at room-temperature, carbogenated 4-(2-hydroxyethyl)-1-piperazineethanesulfonic acid (HEPES) aCSF of the following composition: 92 mM NaCl, 2.5 mM KCl, 1.25 mM NaH2PO4, 30 mM NaHCO3, 20 mM HEPES, 25 mM glucose, 2 mM thiourea, 5 mM Na-ascorbate, 3 mM Na-pyruvate, 2 mM CaCl_2_.4H_2_O and 2 mM MgSO_4_·7H_2_O. Slices were stored for 1-75 hours with minimal submersion (~2 mm depth from the air-liquid interface) on soft nylon netting before transfer to the recording chamber for patch clamp recording. For prolonged slice incubation times beyond 12 hours in the holding chamber, the HEPES aCSF was refreshed every 12 hours and the Brain Slice Keeper-4 holding chamber was thoroughly cleaned to mitigate bacterial growth and slice deterioration. The inclusion of 20 mM HEPES in the aCSF formulations (with concomitant increase in NaHCO_3_) was important to ensure adequate pH buffering and to additionally reduce edema over extended periods of time with the slices in the holding chamber ^22,25^. The osmolality of all solutions was measured at 300-310 mOsmoles/Kg.

### Human ex vivo brain slice cultures

The adult human *ex vivo* brain slice culture method was modified based on existing procedures previously optimized for juvenile rodent hippocampal slice culture and patch clamp electrophysiology ^26^ and adapted to incorporate the NMDG protective recovery method ^22^. Surgical specimens were processed in sterile-filtered and chilled NMDG-aCSF that was saturated with carbogen gas to maintain tissue oxygenation throughout the processing steps. Slices of 300 μm thickness (as compared to 350 μm for acute slices above) were prepared under sterile technique in a biosafety hood using a manual tissue chopper (Leica) equipped with a digital micrometer. Tissue slices were evaluated under a dissecting microscope, and each individual slice was carefully separated and trimmed of any damaged tissue or attached pia. The slices were then transferred into a container of sterile-filtered NMDG aCSF that was pre-warmed to 32-34°C and continuously bubbled with carbogen gas. After 12 min recovery incubation, the slices were transferred onto membrane inserts (PICMORG, Millipore) in 6 well culture plates with 1 mL of slice culture medium underneath, and the residual liquid on the surface of the slices was carefully removed by micropipette. The slice culture medium composition included 8.4 g/L MEM Eagle medium, 20% heat-inactivated horse serum, 30 mM HEPES, 13 mM D-glucose, 15 mM NaHCO_3_, 1 mM ascorbic acid, 2 mM MgSO_4_·7H_2_O, 1 mM CaCl_2_·4H_2_O, 0.5 mM GlutaMAX-I, and 1 mg/L insulin, similar to a slice culture medium previously described for rodent hippocampal slice cultures ^26^. The slice culture medium was carefully adjusted to pH 7.2-7.3, osmolality of 300-310 mOsmoles/Kg by addition of pure H2O, sterile-filtered and stored at 4°C for up to two weeks. Culture plates were placed in a humidified 5% CO2 incubator at 35°C and the slice culture medium was replaced every 2-3 days until end point analysis.

### Patch clamp electrophysiology and live imaging

Human neocortical brain slices were transferred one at a time to the recording chamber of a SliceScope Pro (Scientifica) upright microscope equipped with infrared differential interference contrast (IR-DIC) optics and epifluorescence. The slices were perfused at a rate of 4 ml/min with 32-35°C carbogenated recording aCSF of the following composition: 119 mM NaCl, 2.5 mM KCl, 1.25 mM NaH_2_PO_4_, 24 mM NaHCO_3_, 12.5 mM glucose, 2 mM CaCl_2_·4H_2_O and 2 mM MgSO_4_·7H_2_O. Whole-cell patch-clamp recordings were obtained from visually identified cell bodies of neurons using borosilicate glass pipettes (King Precision Glass; glass type 8250) pulled on a horizontal pipette puller (P1000, Sutter Instruments) to a resistance of 3-5.5 MΩ when filled with the internal solution containing 130 mM K-gluconate, 10 mM HEPES, 0.3 mM EGTA, 4 mM Mg-ATP, 0.3 mM Na_2_-GTP and 2 mM MgCl_2_. The pH was adjusted to 7.3-7.4 with concentrated KOH and the osmolality was adjusted to 290 mOsmoles/Kg using sucrose. In some experiments, the internal solution was supplemented with 50 μM Alexa 594 dye to assist with visualization of cell filling. The theoretical liquid junction potential was calculated at −13 mV and was not corrected. After attaining the whole-cell current clamp mode, pipette capacitance was neutralized and bridge balance applied. All recordings were carried out at 32-34°C with a perfusion rate of ~4 mL/min.

Ca^2+^ imaging was performed on live human *ex vivo* brain slices using widefield epifluorescence illumination or a custom spinning disc confocal microscope (Yokagawa CSUX) attached to the patch clamp rig.

### HSV-1 amplicon vectors and viral transduction

All HSV-1 amplicon vectors were obtained from the Massachusetts Institute of Technology (MIT) Viral Core Facility. The majority of experiments were carried out using stock vectors: HSV-hEF1α-EGFP, HSV-hEF1α-EYFP, HSV-hSyn1-EYFP, HSV-CaMKIIα-EYFP, ST-HSV-ChR2(H134R)-EYFP, HSV-hEF1α-ChR2(H134R)-EYFP. We also created the custom vector HSV-hEF1α-GCaMP6s-P2A-nls-dTomato for this study. The GCaMP6s-P2A-nls-dTomato coding region from Addgene plasmid #51084 was shuttled into the HSV-1 amplicon vector backbone using Gateway cloning. The method of virus packaging and viral vector designs are proprietary to the MIT Viral Core Facility; however, essential aspects of the packaging protocols have been described previously ^27,28^. Functional virus titers were between 1-3 × 10 ^9^ infecting units per mL as measured in PC12 cells. To transduce human slice cultures with HSV-1 amplicons, the concentrated viral stock solutions were directly applied to the slice surface using a fine micropipette at least 1 hour after the time of initial plating such that the slice could first equilibrate. The tip of the pipette was used to carefully spread the virus over the whole slice surface while maintaining surface tension of the viral solution over the slice surface and intentionally avoiding spreading of the solution onto the surrounding membrane insert, to the extent possible.

### Histology and imaging

Brain slices were fixed in 4% paraformaldehyde in phosphate buffered saline (PBS) at 4°C overnight or up to 48 hours and then transferred to PBS with 0.01% sodium azide as a preservative. Fixed slices were thoroughly washed with PBS to remove residual fixative, then blocked for 1 hr at room temperature in PBS containing 5% normal goat serum and 0.2% Triton-X 100. After blocking, slices were incubated overnight at 4°C in blocking buffer containing one or more of the following antibodies: mouse anti-NeuN (1:1000 dilution; Chemicon MAB377), chicken anti-GFP (1:500 dilution; Aves GFP-1020), or rabbit anti-red fluorescent protein (1:500 dilution; Rockland 600-401-379). Following the overnight incubation, slices were washed for 20 min three times with PBS and then incubated for 2 hr at room temperature in Alexa dye-conjugated secondary antibody (1:1000; Invitrogen, Grand Island, NY). Alexa dyes utilized were: Alexa Fluor 488 (goat anti-chicken), Alexa Fluor 555 (goat anti-rabbit and goat anti-mouse), and Alexa Fluor 647 (goat anti-mouse). Slices were washed for 20 min three times with PBS, followed by DAPI nuclear staining for 15 min. Slices were treated with 0.1% Sudan Black B (Sigma-Aldrich, Saint Louis, MO) in 70% ethanol to quench lipofuscin autofluorescence. The slices were then dried on glass microscope slides and mounted with Fluomount G (SouthernBiotech, Birmingham, AL). Slides were stored at room temperature in the dark prior to imaging. Whole slice montage images were acquired with NIS-Elements imaging software on a Nikon Eclipse Ti-E Inverted Microscope System equipped with a motorized st age and epifluorescence illumination. Confocal z-stack images were acquired on an Olympus Fluoview 1000 laser scanning confocal microscope equipped with 488 nm, 543 nm, 594 nm, and 647 nm excitation laser lines.

### Data analysis and statistical testing

Electrophysiological data were analyzed using custom analysis software written in *Igor Pro* (Wavemetrics). All measurements were made from the resting membrane potential. Input resistance (R_N_) was calculated from the linear portion of the current-voltage relationship generated in response to a series of 1s current injections (-150 to +50 pA, 20 or 50 pA steps). The maximum and steady state voltage deflections were used to determine the maximum and steady state R_N_, respectively. Voltage sag was defined as the ratio of maximum to steady-state R_N_. Action potentials (APs) were elicited in response to increasing amplitude, 1_s_ direct current injections. Single APs elicited within 40-50 ms of the start of the current injection were selected for analysis of AP kinetics. AP threshold was defined as the voltage at the time corresponding to when the first derivative of the voltage response first exceeded 20 mV/ms. AP amplitude was measured from threshold to peak, with the half-width measured as the width at half maximal amplitude. Trains of 10±3 APs were selected for spike train analyses. Spike frequency accommodation was defined as the ratio of the last inter-spike interval to the first inter-spike interval. Coefficient of variation was defined as the standard deviation/mean of all inter-spike intervals.

Acute brain slice electrophysiology data evaluating features over time post-slicing was analyzed with linear regression and Pearson’s Correlation. Sample size, p values and r^2^ values are reported. Comparison of acute versus cultured pyramidal neuron or acute versus cultured interneuron intrinsic membrane properties, and comparison of virus infected versus uninfected neuron properties was performed using the unpaired t-test with Welch’s correction. Only neurons exhibiting a resting membrane potential more negative than −50 mV and action potential firing with overshoot exceeding 0 mV were included in the analysis. This criteria was applied uniformly across all experiments. For virus infection electrophysiology experiments, an initial comparison was made between neurons infected with HSV-hEF1α-EYFP and HSV-hSyn1-EYFP. These groups were not statistically different for any parameter examined; thus, we pooled these groups and refer to them collectively as HSV-EYFP infected neurons. For experiments comparing various neuron properties for tumor versus epilepsy cases, the Mann-Whitney U test was applied, with median values and p values reported, where significant.

Analysis of GCaMP6s-related fluorescence transients was performed as follows: Ca^2+^ transients were measured as the relative changes in fluorescence: ΔF/F(t) =(F(t)-F_0_)/(F_o_-F_B_), where F_o_ is baseline fluorescence measured ~200 ms before stimulation and F_B_ is the background fluorescence measured from a section with minimal signal. Fluorescence was calculated in Image J as mean intensity from the perisomatic region of cells of interest.

### Data Availability

The datasets generated during and/or analyzed during the current study are available from the corresponding author on reasonable request.

## Results

### Extended viability of adult human ex vivo brain slices

The generally accepted time window for stable experimentation on rodent acute brain slices (both juvenile and adult) is 6-12 hours *in vitro*, with electrophysiological and structural deterioration observed at later times ^29,30^. In the course of establishing a human *ex vivo* brain slice electrophysiology platform, we observed that adult human *ex vivo* brain slices remained viable far longer than is typical for rodent brain slices prepared under similar conditions. In most cases, adult human neocortical slices remained viable for electrophysiological recordings with incubation in oxygenated aCSF on the bench top throughout the night and into the following day. This was highly desirable given the limited opportunity to obtain living human brain specimens for functional analysis. Thus, we carried out an analysis of human brain slice viability by patch clamp recording of individual neocortical neurons sampled over a wide range of time points *in vitro*. We recorded from 101 pyramidal neurons in neocortical layers 2 and 3 between 1 and 75 hours post slicing. In order to best maintain slice viability with extended incubation times, the aCSF in the holding chamber was refreshed approximately every 12 hours, which was critical and effective to suppress bacterial growth. Viable neurons could be visually identified under IR-DIC optics (~50-100 μm deep in the z dimension) and targeted for recordings throughout the entire 75 hour window. Notably, the sampling was relatively sparse at the later time points (as the majority of brain slices were intentionally utilized for terminal analysis at the earlier times), but nonetheless, the intrinsic and active membrane properties of the recorded neurons were well-preserved across time (Figure 1). Measured electrophysiological features—including action potential (AP) height, AP half-width, AP threshold, sag ratio, input resistance (Rn), and spike frequency accommodation index (SFA)—exhibited very weak or no correlation with time post slicing (all r^2^<0.03, p>0.05). Of note, the resting membrane potential (RMP) exhibited a significant positive correlation with time post-slicing, indicating modest but progressive depolarization during extended incubation times (r^2^=0.11, p<0.001). Related to RMP, the rheobase exhibited a modest negative correlation with time post-slicing (r^2^=0.058, p<0.05), consistent with a progressive but modest neuron depolarization. Time points beyond 75 hours were not investigated, as it became difficult to constrain bacterial growth with brain slices prepared under non-sterile conditions. Nissl staining was performed on 25 μm thick sub-sections of the 5 hr and 72 hr incubated slices, which revealed that the laminar architecture was well-maintained over this time frame, with no overt signs of neuron loss (Supplementary Fig. S1).

**Figure 1:**
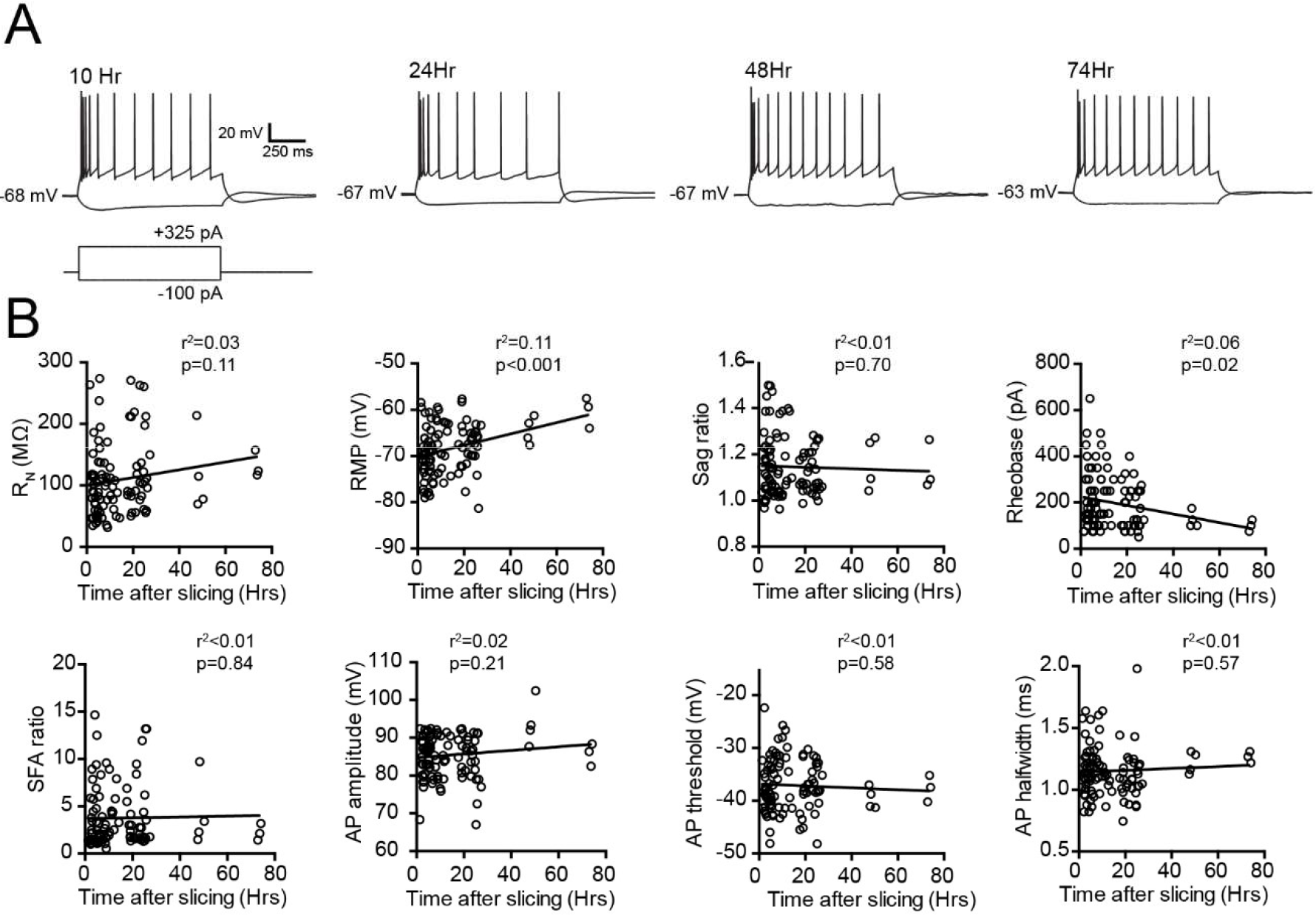
Intrinsic membrane properties of neocortical pyramidal neurons in human *ex vivo* brain slices maintained for extended times *in vitro*. (A) Example traces of membrane voltage responses to hyperpolarizing (-100 pA) and depolarizing (+325 pA) current injection steps at various times postslicing measured from four different neurons. (B) Plots of various electrophysiological features as indicated vs. time post-slicing. Rn, input resistance; RMP, resting membrane potential; SFA, spike frequency adaptation; AP, action potential. n=101 neurons derived from 15 surgical cases (11 epilepsy and 4 tumor).

Given that the surgical specimens included in this study were derived from both tumor and epilepsy patients, we were interested to explore the potential impact of pathology type on neuronal properties. Thus, we performed a sub-analysis of intrinsic membrane properties of neocortical neurons in acute brain slices derived from tumor patients versus epilepsy patients (Supplementary Fig. S2). We focused on the 0-12 hr time period following brain slicing, since this was the time frame at which we had the greatest sampling for both conditions (and noting that sampling of neurons from tumor cases is sparser overall, and especially at longer time points, due to fewer specimens obtained in this study). The median rheobase values were 150.0 pA and 212.5 pA for tumor vs epilepsy cases, respectively, and the distributions in the two groups differed significantly (Mann-Whitney *U* = 214; n=14 neurons from tumor cases and n=44 neurons from epilepsy cases; P < 0.05). The distribution of measured values for all other membrane properties (RN, RMP, Sag, AP amplitude, AP threshold, AP half-width, and SFA) were not significantly different between tumor and epilepsy cases (n=14 neurons from tumor cases and n=44 neurons from epilepsy cases; P > 0.05).

### An adult human brain slice culture platform suitable for rapid genetic labeling

The robustness and longevity exhibited by these human *ex vivo* brain slices lends naturally to organotypic brain slice culture applications. We adapted a rodent hippocampal brain slice culture protocol to our adult human neocortical brain slices ^26^ (see materials and methods for details). Under these conditions, adult human *ex vivo* brain slices could be maintained for at least 1 week *in vitro* (the longest time point examined in this study) in a 5% CO_2_ incubator at 35°C. Histological examination of the cultured brain slices revealed that the laminar architecture of the neocortex remains intact, with no overt signs of cell dispersion (Figure 2a). In addition, neuronal density at 2 days *in vitro* (DIV), as judged by immunohistochemical (IHC) staining for NeuN, was well-preserved and comparable to acute slices from the same surgical specimen (Figure 2b-d).

**Figure 2:**
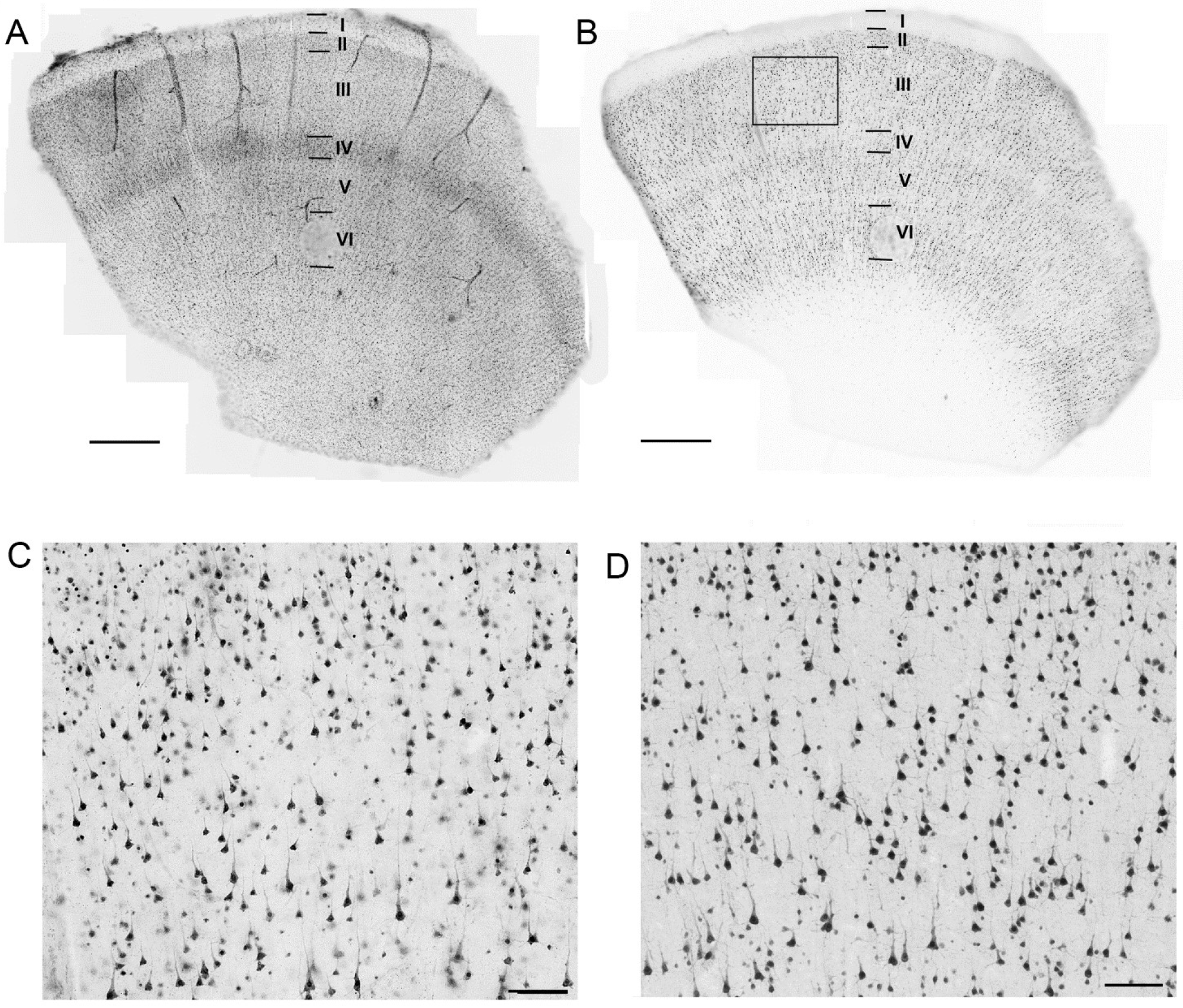
Human neocortical brain slices maintain laminar architecture and neuronal integrity during the early culture period. Neocortical slice culture at 2 days *in vitro* labeled with DAPI (A) or anti-NeuN (B). (C) Higher magnification of boxed region in (B). (D) Anti-NeuN labeling for the matched region of an acute brain slice from the same surgical specimen. Scale bars: 1 mm (A, B) and 100 μm (C,D).

Quantitative descriptions of the physiological properties of adult human neurons in brain slice cultures are necessary to establish the functional integrity of the platform. To directly address this point, we performed whole-cell patch clamp recordings on pyramidal neurons and interneurons in human brain slice cultures at 2-3 DIV (Figure 3a,b). Intrinsic membrane properties were compared to pyramidal neurons or interneurons recorded in acute (<24 hr post slicing) brain slices (Figure 3c). We observed three properties of human pyramidal neurons that were altered in 2-3 DIV slice cultures relative to acute slices, including RMP (culture: mean= −62.0 ± 2.0 mV, n=10 neurons; acute: mean= −68.9 ± 0.6 mV, n=82 neurons; **p < 0.01), AP amplitude (culture: mean= 70.8 ± 3.4 mV, n=10 neurons; acute: mean= 85.4 ± 0.6 mV, n=82 neurons; **p < 0.01), and AP halfwidth (culture: mean= 0.84 ± 0.04 msec, n=10 neurons; acute: mean= 1.15 ± 0.02 msec, n=82 neurons; ***p < 0.001). Other features examined, including Rn, sag ratio, and AP threshold were not significantly altered (p > 0.05 for all values). In contrast to pyramidal neurons in culture, interneurons did not exhibit altered intrinsic properties across all parameters measured, including Rn, RMP, Sag ratio, AP amplitude, AP threshold, and AP half-width, as compared to interneurons from acute slices.

**Figure 3:**
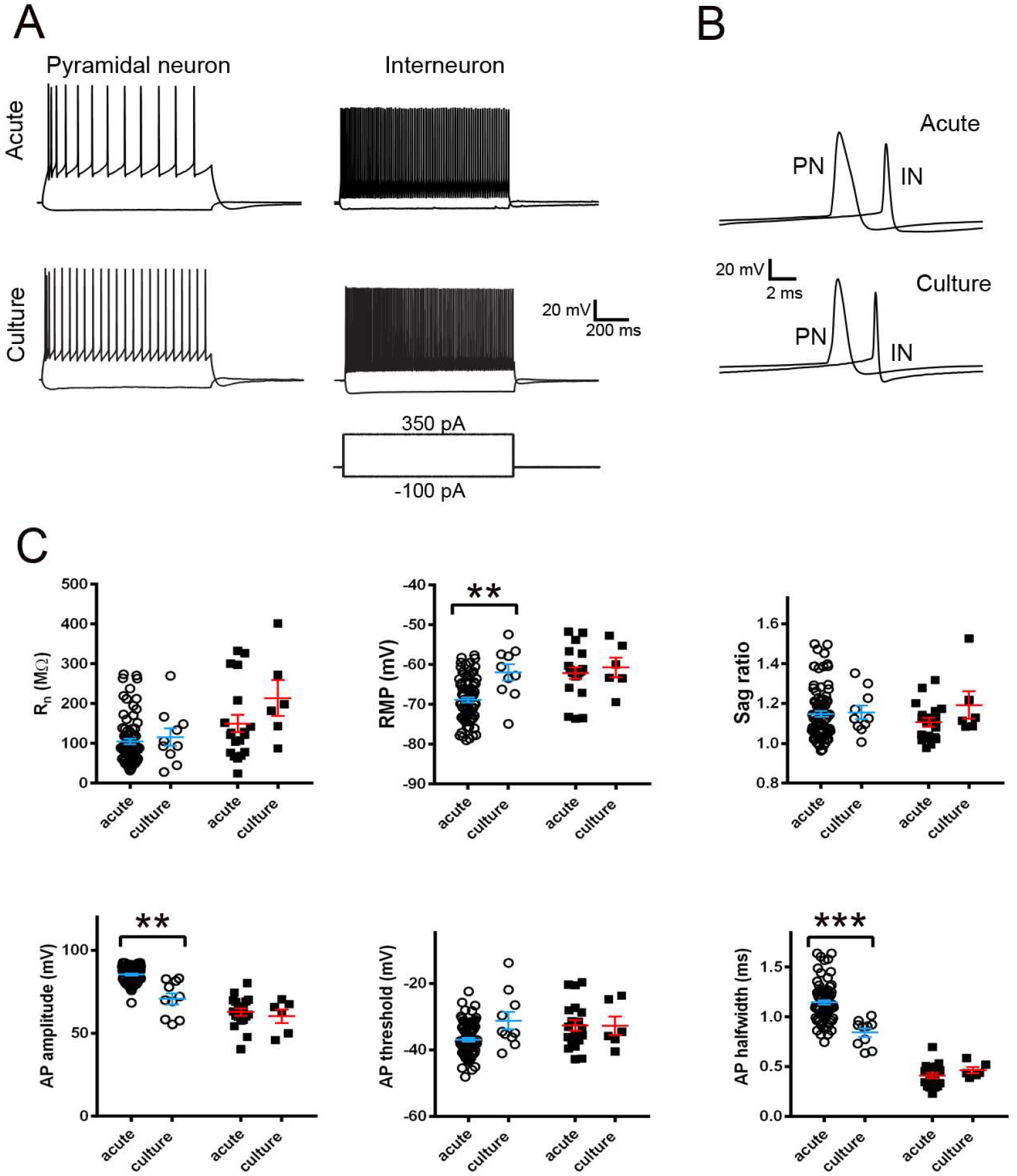
Intrinsic membrane properties for pyramidal neurons and interneurons recorded in acute versus cultured human *ex vivo* brain slices. (A) Example traces of acute versus culture pyramidal neuron (left) and interneuron (right) action potential firing in response to 1s current injection steps. (B) Expanded view of single action potential waveforms (PN, pyramidal neuron; IN, interneuron). (C) Comparison of acute versus cultured slice intrinsic membrane properties for pyramidal neurons (open circles, blue bars) and interneurons (black squares, red bars). Pyramidal neurons: acute (<24 hr post slicing), n=82 neurons; culture (2-3 DIV/DPI), n=10 neurons; Interneurons: acute (<24 hr post slicing), n=19 neurons; culture (2-3 DIV/DPI) n=6 neurons. * p < 0.05, ** p < 0.01, *** p < 0.001, unpaired t-test with Welch’s correction.

We next explored the feasibility of genetic labeling in our culture system. We chose to focus on HSV-amplicons, a double-stranded enveloped DNA virus type with a well-described rapid onset of transgene expression and strong neuronal tropism ^23,31^. HSV-1 amplicon vectors encoding EYFP under the human Synapsin-1 (hSyn1), human elongation factor 1α (hEF1α), or rat Calcium/calmodulin-dependent protein kinase type II alpha (CaMKIIα) promoter fragments were applied directly to the surface of the slices on the day of culturing. Viral transduction efficiency and reporter transgene expression was assayed by epifluorescence microscopy at various time points following virus infection. Faint native fluorescence could be detected in less than 24 hours post-infection, whereas peak expression was observed by 2-3 days post infection (DPI), followed by a gradual decline in transgene expression levels at later times (Supplementary Fig. S3; see also Supplementary Fig. S4). Widespread viral transduction and transgene expression was achieved, with labeled neurons located across the cortical depth, as visualized by EYFP native fluorescence and immunohistochemical (IHC) detection of GFP protein (Figure 4). Although virus infected neurons could be observed throughout the z-depth of the slice, a gradient of labeling density was typically observed with more transgene expressing neurons nearest the exposed slice surface.

**Figure 4:**
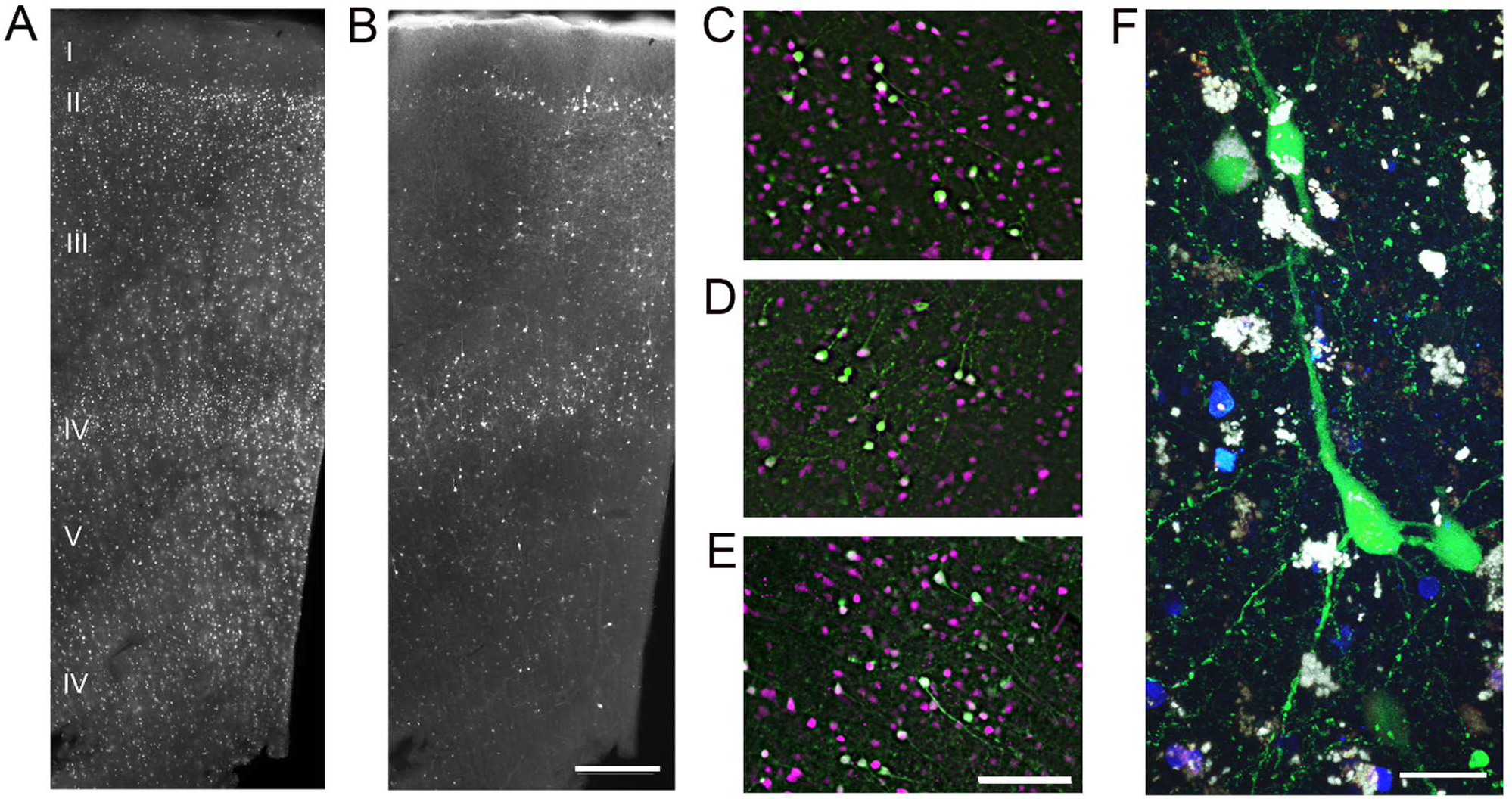
Rapid viral genetic labeling of neocortical neurons in human *ex vivo* brain slice cultures. (A) anti-NeuN staining throughout the cortical depth. (B) Anti-GFP staining in the same field of view as in (A) at 2 DIV/2DPI with HSV-CaMKIIα-EYFP. Scale bar: 300 μm. (C-E) Co-immunostaining with anti-GFP (green) and anti-NeuN (magenta) for human slice cultures infected with HSV-CaMKIIα-EYFP (C), HSV-hSyn1-EYFP (D), and HSV-hEF1α-EGFP (E) at 2DIV/2DPI demonstrated restricted neuronal expression. Scale bar: 100 μm(F) High magnification live confocal image of native fluorescence for a cultured human slice infected with HSV-hEF1α-EGFP at 3 DIV/3DPI. Nuclei are shown in blue and lipofuscin granules appear as white. Scale bar: 15 μm.

Although virus-labeled neurons were found throughout all cortical layers, there was a reproducible bias with the most extensive transduction of neurons within layer 2, deep layer 3, and layer 4 (Figure 4b). Co-immunostaining for NeuN and GFP protein revealed that the viral transduction was exclusively neuronal in human neocortical brain slice cultures, with three distinct HSV-1 amplicon vectors tested (Figure 4c-e; GFP+/NeuN+: HSV-hEF1α-EYFP, 75/75; HSV-hSyn1-EYFP, 59/59; HSV-CaMKIIα-EYFP, 61/61). Both neuron-specific (hSyn1 and CaMKIIα) and ubiquitous (hEF1α) promoters yielded pure neuronal expression in the context of HSV-1, indicating that the viral tropism of HSV-1 is exclusive for neuronal types in the brain, as previously reported ^31^.

The prospective marking of neurons with fluorescent reporters in adult human neocortical slices can be useful for targeted functional analysis using patch clamp electrophysiology. This approach is also very valuable for assessing functional changes related to viral transduction. We carried out whole-cell patch clamp recordings to compare passive and active membrane properties of HSV-infected neocortical pyramidal neurons and neighboring uninfected pyramidal neurons between 2-3 DIV/DPI, which represents an early time window that is well matched to peak virus-mediated transgene expression. HSV-EYFP infected neurons were easily visualized and targeted for patch clamp recordings by detection of bright native fluorescence at the somata. Virus-infected and uninfected neurons exhibited comparable passive membrane properties (Figure 5), with no significant differences observed for RMP (HSV-EYFP: mean = −64.0 +/- 1.5 mV, n=18; uninfected: mean = −62.1 +/- 1.9 mV, n=10; p=0.44) or Rn (HSV-EYFP: mean = 159.2 +/- 22.7 MOhms, n=18; uninfected: mean = 118.1 +/- 22.3 MOhms, n=10; p=0.24). In addition, there were no significant differences in active membrane properties, including AP amplitude (HSV-EYFP: mean = 62.8 +/- 3.2 mV, n=18; uninfected: mean = 68.5 +/- 3.8 mV, n=10; p=0.27), AP threshold (HSV-EYFP: mean = −28.9 +/- 1.8 mV, n=18; uninfected: mean = −30.5 +/- 2.5 mV, n=10; p=0.60), or AP half-width (HSV-EYFP: mean = 0.91 +/- 0.04 ms, n=18; uninfected: mean = 0.82 +/- 0.045 mV, n=10; p=0.14). Sag ratio was the sole parameter examined that was slightly but significantly different between groups (HSV-EYFP: mean = 1.07 +/- 0.02, n=18; uninfected: mean = 1.15 +/- 0.03, n=10; p<0.05).

**Figure 5:**
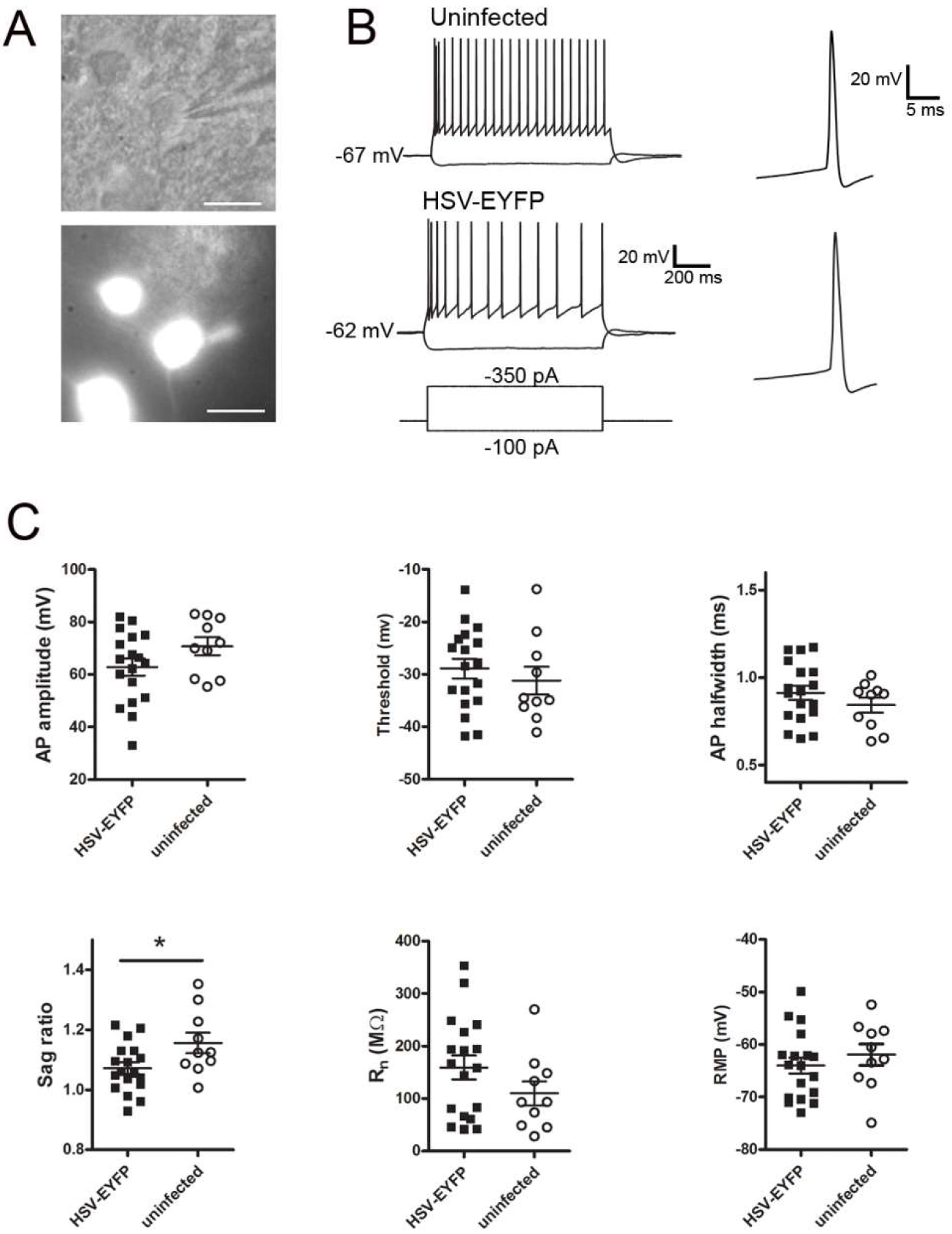
Intrinsic membrane properties of HSV-infected versus uninfected neocortical pyramidal neurons in human *ex vivo* brain slice cultures. (A) IR-DIC (top) and epi-fluorescent (bottom) image of a recorded HSV-EYFP infected pyramidal neuron. Scale bar: 20 μm. (B) Example voltage responses to hyperpolarizing and supra-threshold depolarizing current injections for an uninfected (top) and HSV-EYFP infected (bottom) pyramidal neuron. Examples of single APs are shown at a faster time scale at right. (C) With the exception of voltage sag ratio, the properties of HSV-EYFP infected (black squares, n=18 neurons) and uninfected (open circles, n=10) cortical pyramidal neurons were comparable for recordings carried out between 2-3 DPI/DIV. * p < 0.05, unpaired t-test. These data were derived from 8 surgical cases (7 epilepsy and 1 tumor case).

### Functional analysis of human neocortical neurons using genetically-encoded probes

We next investigated viral expression of the optogenetic actuator ChR2 ^32^ for optical control of neuron firing in adult human brain slice cultures. Slices were infected with HSV-ChR2-EYFP and patch clamp recordings were carried out over the first 72 hours post-infection to determine the time point at which the ChR2-EYFP expression reached sufficient levels to drive action potential firing with blue light. The light-evoked peak photocurrent amplitude increased sharply from 185.1 pA +/- 10.8 pA at 1 DPI (n=4 neurons) to 796.1 pA +/-74.8 pA at 2 DPI (n=6 neurons; values are reported as mean +/- standard error of the mean), but remained relatively high at 3 DPI (mean = 712.5 pA +/- 95.4 pA; n=3), (Supplementary Fig. S4). The ChR2-EYFP expression level at 2-3 DPI was sufficient to optically induce reliable action potential firing over a range of stimulation frequencies using 3 ms blue light pulses in a subset of neurons. The highest expressing neurons exhibited robust peak photocurrent amplitudes approaching (but rarely exceeding) 1 nA. Faithful entrainment of action potential firing was achieved up to 10 Hz. Functional ChR2-EYFP was observed in both pyramidal neurons and interneurons (data not shown).

We also explored HSV-mediated expression of the genetically-encoded calcium indicator GCaMP6s ^33^ for optical monitoring of activity in human brain slice cultures. The overall extent of viral transduction with HSV-GCaMP6s was assessed by examining the distribution of nuclear-localized dTomato reporter. The multi-cistronic viral vector design with a constitutive red reporter facilitated the identification of healthy GCaMP6s-expressing neurons that exhibited low basal activity. Live confocal imaging was performed at 4DIV/4DPI to capture spontaneous functional activation of GCAMP6s expressing neurons observed as transient increases in cytosolic green fluorescence over baseline fluorescence (Figure 6a,b). The majority of virus-infected neurons were quiescent and exhibited low basal green fluorescence, which was verified by the presence of bright nuclear dTomato fluorescence but very low cytosolic green fluorescence. Similar observations of spontaneous activation of HSV-GCaMP6s transduced human neurons was replicated at various time points between 3-6 DIV/DPI and across multiple virus-infected slices from 5 distinct human slice culture preparations. To show that the observed Ca^2+^ transients reflected action potential firing of the neurons, we performed simultaneous epifluorescence imaging and patch clamp recording and electrical stimulation via the patch pipette to drive neuron firing at frequencies from 1-50 Hz. Graded responses in the amplitudes of the peak fluorescence were observed, as expected, with a detection threshold of ~3 action potentials (Figure 6c). In a separate experiment, we transduced cultured human hippocampal slice cultures derived from a temporal lobe epilepsy patient with HSV-GCaMP6s virus and performed live confocal imaging of dentate gyrus granule cells at 4 DIV/4DPI (Supplementary Movie S1). Robust carbachol-evoked Ca^2+^ transients were detected across a population of neurons in the granule cell layer (Figure 6d). Collectively, these experiments demonstrate that GCaMP6s-expressing neurons in adult human brain slice cultures were functionally intact and amenable to monitoring of spontaneous or evoked activity over time.

**Figure 6:**
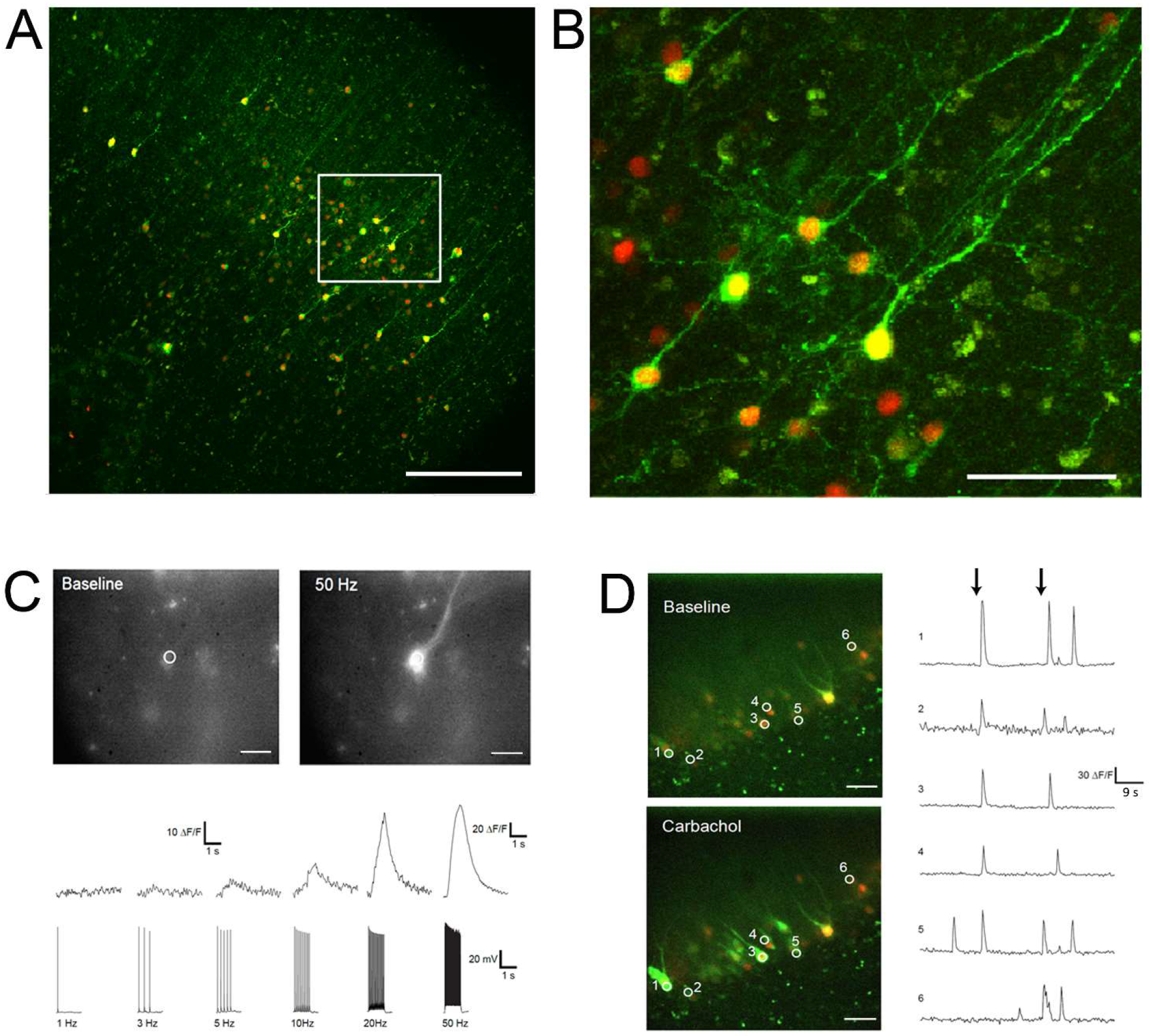
Rapid viral expression of the genetically-encoded Ca^2+^ indicator GCaMP6s in human *ex vivo* brain slice cultures for optical monitoring of neuronal activity. (A) Representative confocal z-stack image of an HSV-hEF1α-GCaMP6s-P2A-nls-dTomato infected human neocortical brain slice culture at 4DIV/4DPI. Constitutive expression of nuclear dTomato (red) serves as a reporter of virus-infected neurons, whereas active neurons exhibit additional cytoplasmic fluorescence of GCaMP6s (green). Scale bar: 300 μm (B) Higher magnification view of the boxed region in (A). Scale bar: 100 μm. (C) Fluorescence imaging of baseline and electrical stimulus-evoked Ca^2+^ transients in an HSV-hEF1α-GCaMP6s-P2A-nls-dTomato infected human neocortical pyramidal neuron at 4 DIV/DPI (top panels). The region of interest (ROI) for analysis is indicated by a white circle. Scale bar: 30 μm. Recorded Ca^2+^ transients (ΔF/F) aligned to corresponding patch clamp electrode-evoked action potential firing over 1 sec bouts at the indicated stimulation frequencies (bottom panel). Note the different (ΔF/F) scale for the 50 Hz Ca^2+^ transient. (D) Dual-color, spinning disc confocal population imaging of the granule cell layer in HSV-hEF1α-GCaMP6s-P2A-nls-dTomato infected human *ex vivo* hippocampal slices at 4 DIV/DPI. Scale bar: 40 μm. Single plane images are shown for baseline fluorescence and the subsequent fluorescence increase following bath application of 100 μm carbachol (left panels). The 6 selected ROIs are indicated by numbered white circles. Line plots showing the carbachol evoked Ca^2+^ transients for each individual ROI. Note the strong, largely synchronous population response with carbachol stimulation (black arrows indicate time of bath application). These data were derived from three different surgical cases.

## Discussion

Earlier studies using adult human *ex vivo* brain slices alluded to an extended viability of the tissue but with no direct experimental evidence in support of this claim ^6,8,34,35^. It seems this anecdotal evidence has gone largely unnoticed, and perhaps much more is possible in the way of human brain slice functional studies than has been widely appreciated in the neuroscience community. We provide the first detailed functional characterization to substantiate the remarkable viability and longevity of human *ex vivo* brain slices, consisting of a quantitative time course analysis of single neuron electrophysiological properties sampled over 75 hours following removal from the intact brain of a patient and processing into tissue slices. Surprisingly, the neuronal membrane properties for the sampled neurons were well-preserved, with only resting membrane potential and rheobase modestly altered over time. This evidence clearly demonstrates that, under certain conditions, neurons in these brain slices can remain viable and healthy, and that basic functional properties can be continuously measured for extended periods. This is a highly desirable outcome given the limited opportunity to carry out research studies on precious human neurosurgical specimens. The exact procedure and conditions for collection and processing of the surgical specimens are likely to be important to achieving the extended human brain slice viability in our experiments. In particular, we applied the NMDG protective recovery method of brain slice preparation ^22^, a method that improves the viability of adult rodent brain slices but that had not previously been applied to human brain tissue. Apart from the methodological considerations, the exact cellular mechanisms responsible for the remarkable viability of adult human *ex vivo* brain slices (in comparison to rodent brain slices) is of great interest and remains to be explored.

The robustness of our human *ex vivo* brain slices was leveraged to establish an organotypic brain slice culture platform. We opted to focus on cultures of less than one week, where structural and functional properties are reasonably maintained with minimal perturbation, to be as comparable as possible to measurements obtained using gold-standard acute slice preparations. This is in contrast to recent work that explored optimal conditions for long-term cultures of human *ex vivo* brain slices for up to one month *in vitro*, with the earliest time point systematically examined being six days *in vitro* ^20^. This extended culture time was rationally posited as necessary for viral genetic labeling and perturbation experiments, with direct feasibility emerging shortly thereafter by a separate group that used lentiviral expression of ChR2-EYFP in human *ex vivo* slice cultures maintained for a minimum of two weeks *in vitro* ^21^. ChR2-EYFP expression levels sufficient to optically drive neuron firing and a pharmacologically-inferred synaptic transmission were achieved in this longer expression period. Despite the importance of this technological breakthrough, the structural and functional integrity of these virus-infected slice cultures are unclear due to the limited evidence provided. Furthermore, the morphology of the virus-infected neurons was suggestive of potential culture artifacts. These issues may constrain the overall utility of such extended cultures for deriving meaningful properties of human neurons and local circuits.

In the present work, we circumvent some of the confounding effects of prolonged culture by the judicious choice of virus type for rapid onset and high-level transgene expression. HSV-1 amplicon vectors exhibited the following desirable features: strong neurotropism, rapid onset of transgene expression, high peak expression levels, and minimal neurotoxicity in this early time window ^23,31^. We achieved widespread labeling of human neocortical neurons in cultured brain slices within days following transduction with HSV-1 amplicon vectors encoding fluorescent reporters and optogenetic probes. This viral labeling is considerably more rapid than the two weeks required to achieve functional expression of ChR2-EYFP in human slices using lentivirus ^21^. Despite this much shorter period following viral transduction, we achieved equivalent levels of ChR2-mediated photocurrent in human pyramidal neurons, as well as precise light-evoked action potential firing. Furthermore, we provide the first demonstration and feasibility of applying a genetically-encoded calcium indicator GCaMP6s to monitor the activity of adult human neurons, including analysis of population dynamics, in local brain circuits. By combining optogenetic actuators and spectrally-compatible activity indicators, it should be possible to carry out all-optical interrogation of human brain circuits in this brain slice platform, similar to what has been demonstrated in the mouse brain ^36,37^.

The ability to deliver viral payloads to particular cell types in adult human brain tissue, combined with a diverse suite of modern molecular genetic tools, could accelerate human brain research with particular emphasis on cellular resolution functional studies. Although our current study builds towards this aim, further innovations are necessary to both refine and diversify the genetic expression strategies that are applicable to human *ex vivo* slices. Previous work has demonstrated the feasibility of achieving cell type-specific transduction and transgene expression with HSV-1 vectors in rodent brain ^38–41^, including for broad classes of neocortical neurons ^42^. It is also anticipated that viral vectors utilizing Cre recombinase-dependent expression strategies can be applied, and multiple studies have succeeded at applying intersectional expression strategies using viral vectors in the non-human primate brain ^43,44^. Whether such viral tools can achieve high specificity with sufficiently rapid transgene expression in the context of human *ex vivo* brain slice cultures is still an open question. This represents an important next step towards enabling a precise genetic dissection of human local neocortical circuits in both health and disease. This platform may also serve as a valuable and unique new proving ground for viral vectors on the path toward human gene therapy applications. Such a strategy could help to minimize negative impacts to patients by first verifying tolerability and functionality of the viral vectors directly on adult human brain tissue *in vitro*, and for confirming selective targeting of cell types of interest. Taken together, the overall innovative approach of rapid genetic labeling in adult human *ex vivo* brain slices may be suitable for addressing a wide range of experimental questions about the functions of human brain cell types and circuits that were previously not feasible.

## Author Contributions

J.T.T., E.L., and C.K. conceived and designed the study. J.T.T. developed the human *ex vivo* brain slice culture and viral transduction platform. J.T.T., B.K., and P.C. performed the electrophysiological recording experiments. J.T.T. and R.D.F. conducted the histological staining and imaging experiments. J.T.T. and B.K. analyzed the data and J.T.T., B.K., wrote the manuscript with input from E.L., and C.K.. J.G.O., A.L.K., R.P.G., and C.C. provided human neurosurgical specimens deemed suitable for research purposes. C.D.K. provided neuropathology expertise and consultation and supported collection and transport of surgical specimens at Harborview Medical Center. R.G.E. provided support in establishing the neurosurgeon network at the University of Washington Hospital and Harborview Medical Center sites.

## Acknowledgements

We wish to thank Dr. Gábor Tamás and colleagues for generous help with establishing the acute human *ex vivo* brain slice platform. We thank the Allen Institute for Brain Science Tissue Procurement and Processing teams (especially Nick Dee and Julie Nyhus) and the Facilities teams for help in coordinating logistics of human surgical tissue collection, transport and processing. We are also grateful to our collaborators at the local hospital sites, including Tracie Granger, Caryl Tongco, Matt Ormond, Jae-Guen Yoon, Nathan Hansen, Niki Ellington, Rachel Iverson (Swedish Medical Center), Carolyn Bea, Gina DeNoble, and Allison Beller (Harborview Medical Center). The HSV-1 amplicon vectors were packaged and distributed by Dr. Rachael Neve from the Viral Core at the Massachusetts Institute of Technology (now at MacLean Hospital, Boston, MA). A special thanks to Dr. Andres Barria (University of Washington Department of Physiology & Biophysics) for help with brain slice culturing techniques. We are grateful to our Allen Institute colleagues Hongkui Zeng, Boaz Levi, John Mich, and Jim Berg for critical reading and advice on the manuscript, Luke Campagnola for help with GCaMP data analysis, and Susan Sunkin for help with project management. We wish to thank Paul G. Allen for his vision, encouragement, and support.

